# The Association Between SLIT2 in Human Vitreous Humor and Plasma and Neurocognitive Test Scores

**DOI:** 10.1101/2025.05.03.652072

**Authors:** Sara I. Shoushtari, Easton Liaw, Sreevardhan Alluri, Zahra Sheikh, Sudhir Kumar, Courtney Huynh, Insa M. Schmidt, Steven Ness, Xuejing Chen, Nicole H. Siegel, Sushrut S. Waikar, Thor D. Stein, Weining Lu, Manju L. Subramanian

## Abstract

**Background:** Slit Guidance Ligand 2 (SLIT2) binds Roundabout (ROBO) guidance receptors to direct axon pathfinding and neuron migration during nervous system development. SLIT2 expression has previously been linked to dementia risk.

**Objective:** To study the association between SLIT2 expression in human vitreous humor and plasma samples and neurocognitive test scores in a cross-sectional cohort study utilizing a novel, highly-sensitive Meso Scale Discovery (MSD) assay for SLIT2 detection.

**Methods:** Seventy-nine individuals with a mean age of 55.79 ± 12.03 years underwent eye surgery with collection of vitreous humor, blood (plasma) collection, and neurocognitive assessment. Vitreous humor and plasma samples were analyzed by SLIT2 MSD electrochemiluminescence immunoassay. Associations between SLIT2 levels in vitreous humor and plasma were analyzed using GraphPad Prism.

**Results:** We found up to a 7-fold higher level of SLIT2 in human vitreous humor compared to plasma. Lower vitreous SLIT2 levels were associated with a lower Montreal Cognitive Assessment (MoCA) score and Immediate Recall Verbatim (IRV) z-score, and higher plasma SLIT2 was associated with a lower MoCA score. In multivariate analysis using single and multiple predictor models, the same significant associations were found when adjusted for age, sex, race, diabetic status, diabetic retinopathy status, glaucoma status, and Apolipoprotein E (APOE) genotype.

**Conclusions:** SLIT2 protein levels are significantly associated with MoCA score and IRV z-score in middle-aged individuals. The relationship remained significant when adjusted for demographics, co-morbidity, and APOE genotype, suggesting SLIT2 may be a sensitive biomarker for detection of mild cognitive impairment and early dementia, and warrants further studies.

## BACKGROUND

Neurodegenerative disorders can be a cause of neurocognitive impairment and are classified by pathological changes with potential destruction of neural tissue.^1–3^ One change known in neurodegenerative disorders is an accumulation of proteins causing pathological damage.^4,5^ Protein abnormalities that lead to inadequate clearance and subsequent aggregation can cause this pathological degeneration.^3,6^ These changes can occur decades before the onset of cognitive symptoms.

Mild neurocognitive impairment (MCI) has an approximate prevalence of 6% in individuals aged 60-64 and increases to 25% in individuals aged 80-84.^7^ Approximately 5-10% of individuals with MCI progress to major neurocognitive disorder or dementia each year,^7,8^ the most common of which is Alzheimer’s dementia,^9^ with an estimated prevalence of 6.5% in middle-aged and senior populations, or 5.7 million in the United States.^10^ The prevalence of Alzheimer’s disease is expected to double in the next four decades.^10^ Early-onset dementia is a subset making up approximately 4-6% of Alzheimer’s Disease cases, with an onset before age 65, and is notable for its significant genetic predisposition and rapid progression.^11,12^

Neural-derived proteins have been demonstrated previously to have potential relationships with neurocognitive dysfunction.^13^ For example, increased neurofilament light (NfL) protein levels have been associated with cognitive impairment.^14,15^ Studies have been conducted to assess a variety of proteins involved in the pathological changes of neurodegeneration, such as beta-amyloid (Aβ), t-tau, p-T181-tau, and GFAP.^16,17^ In early-onset dementia, studies have found decreased Aβ and increased total tau in CSF samples.^11^ Our group has studied protein biomarkers in ocular fluid and pathological proteins associated with late-onset Alzheimer’s Disease and MCI.^18–22^ Additionally, we have demonstrated that relatively higher concentrations of neurodegenerative biomarkers exist in ocular fluid, such as vitreous fluid, as compared with plasma.^18^

Slit Guidance Ligand 2 (SLIT2) protein is a ligand that binds to Roundabout (ROBO) guidance receptors expressed during vascular and neuronal development.^23–25^ It is commonly found in cerebral tissue^26,27^ and functions as a chemorepulsive guidance to direct axon pathfinding and neuron migration,^25,28,29^ such as the development of the optic chiasm^30^ and neurodegenerative disease.^31–33^ Proteomic studies have found that SLIT2 levels are increased in Aβ overexpression mouse models^34^ and are correlated with Aβ levels in human tissue.^35^ SLIT2 has also been shown to co-localize with Aβ.^34^ We therefore hypothesize that SLIT2 may also be an additional neurodegenerative biomarker associated with cognitive impairment.

Prior reports^33,35–37^ have suggested a link between SLIT2 protein levels and late-onset dementia and Alzheimer’s Disease through studies on cognitive stimulation in the workplace, proteomic profiling, and forward genetic screening. In a population-based proteomic study, increased levels of plasma SLIT2 protein were shown to be associated with an increased hazard ratio for late-onset dementia, suggesting that SLIT2 may be a potential biomarker candidate for cognitive functional decline and progression to dementia.^36^ However, this proteomic finding has not been validated by a current commercially available SLIT2 immunoassay due to their low sensitivity and dynamic range. Additionally, there is no currently published data on SLIT2 protein levels in the early-onset dementia population.

SLIT2 protein is highly expressed in ocular neurons.^38^ However, SLIT2 in the vitreous humor and its association with cognitive status have not been previously investigated. In this study, we hypothesize that the SLIT2 protein measured in human vitreous humor and plasma is associated with cognitive function based on neuropsychological assessment.

## METHODS

### Study Design

This is a cross-sectional cohort study conducted at Boston University Medical Campus and Boston Medical Center (BUMC/BMC) to assess the association of biomarkers in eye fluid and plasma with neurocognitive test scores. This study utilized previously stored eye fluid and plasma samples from prior research that investigated the association of biomarkers to eye disease and neurocognitive function. The BUMC/BMC institutional review board provided approval and oversight (study reference number H-37370, principal investigator MLS). The study was completed in accordance with institutional ethical standards and the Declaration of Helsinki.

Study participant inclusion criteria included age 18 years or older, primary language English or Spanish, and those scheduled for pars plana vitrectomy (PPV) in at least one eye for a clinical eye condition. Surgical indications for vitrectomy included eye diseases such as rhegmatogenous retinal detachment, macular hole, epiretinal membrane, vitreous hemorrhage, and tractional retinal detachment. Study activities for Spanish-speaking patients were completed with a bilingual research assistant. Written informed consent was obtained from all patients who participated in the study, and participants were not excluded for existing ocular or medical comorbidities. Upon completing the consent process, the study activities included neuropsychological testing, vitreous humor samples collected during PPV, and blood samples collected during a research visit.

### Historical Data Collection

Demographic information and clinical history of enrolled participants were collected from participant-reported questionnaires and electronic medical records. Demographic information included age, race, and sex. Clinical history included past medical history and ophthalmic history. A baseline ophthalmic exam, including assessment of common eye conditions such as glaucoma, age-related macular degeneration, and diabetic retinopathy, was completed prior to eye surgery.

### Neuropsychological testing

Neuropsychological testing was conducted with all enrolled participants within a 3-month window before or after their surgery date. Our group collaborated with the Boston University Alzheimer’s Disease and Research Center, which recommended neuropsychological testing within 180 days of fluid specimen collection and participated in training research assistants to conduct the assessments. Validated tools for testing included the Montreal Cognitive Assessment (MoCA)^39^ and the Immediate Recall Verbatim (IRV).^40^ Test scores (MoCA raw score and IRV z-score) are standardized and allow for comparison between individuals.^41,42^

### Biospecimen Collection

Vitreous and blood samples were collected between September 2019 and August 2021. Vitreous samples were collected at the start of each vitrectomy procedure by the surgical team. 0.5–1.0 mL of undiluted vitreous fluid was aspirated via the vitrectomy probe into an attached sterile 3-mL syringe. Infusion of saline into the vitreous cavity was immediately undertaken in order to re-pressurize the posterior chamber of the eye, and the vitrectomy subsequently proceeded according to the standard of care for the respective ocular condition. The syringe containing the vitreous specimen was capped using sterile technique and directly handed to a research assistant, who labeled it with a predetermined non-identifiable study number and placed the sample on ice for transport. Blood samples were collected from all study participants at a research visit prior to surgery, and eighteen milliliters of whole blood were drawn into Ethylenediaminetetraacetic acid (EDTA)-treated purple top tubes and also placed on ice for transport. Once collected, vitreous and blood biospecimens were immediately centrifuged, aliquoted into 200 microliter Eppendorf tubes at the Molecular Genetics Core Laboratory (MGCL) at BU Medical Campus and stored in −80°C freezers. De-identified blood samples were processed into their component plasma and buffy coat. Samples were thawed once prior to immunoassay analysis between August and November 2023, with no additional freeze-thaw cycles. Plasma and vitreous samples were utilized for SLIT2 protein analysis, and the buffy coat was used for Apolipoprotein E (APOE) genotyping.

### SLIT2 MSD Assay to measure protein expression in the vitreous and plasma

To measure the low SLIT2 protein concentration in the human vitreous humor and plasma samples, we developed a highly sensitive Meso Scale Discovery (MSD) SLIT2 immunoassay using a SLIT2 antibody pair (Catalog # ab253658, Abcam, Waltham, MA, USA) and GOLD 96-Well Small Spot Streptavidin SECTOR Plates (Catalog # L45SA, Meso Scale Diagnostics, Rockville, MD, USA) with an electrochemiluminescence detection technique, which has a greater dynamic range and more precise and accurate readings for low concentrations of SLIT2 at picogram levels. Details on this internally developed SLIT2 MSD assay are currently in preparation and will be reported separately. In a process similar to a sandwich enzyme-linked immunoabsorbent assay (ELISA), we detected and quantified standard SLIT2 recombinant protein and unknown SLIT2 concentration in human vitreous humor and plasma samples using the MESO QuickPlex SQ120 instrument (Catalog # AI1AA-0, Meso Scale Diagnostics, Rockville, MD, USA).

### APOE Genotyping

APOE is a major susceptibility gene, and relative allele copies can confer a higher risk of Alzheimer’s;^43–45^ therefore, APOE genotyping is sometimes used as a clinical risk evaluation tool.

To determine the APOE genotype, DNA was extracted from the buffy coat, and single nucleotide polymorphisms from the APOE gene (National Center for Biotechnology Information SNPs rs429358 and rs7412) were examined using TaqMan assays (Applied Biosystems, Foster City, CA).

### Statistical Analysis

Statistical analyses were performed using GraphPad Prism (Version 10.2.2 341) for MacOS, GraphPad Software, Boston, MA, USA, www.graphpad.com). A p-value of less than 0.05 was considered statistically significant.

SLIT2 protein levels and neurocognitive assessment score association testing, using neurocognitive scores as a continuous variable, was performed using univariate and multivariate linear regression. Univariate analysis included both SLIT2 protein plasma and vitreous levels as predictors of MoCA raw score and IRV z-score. Additionally, analysis was performed with participants grouped based on MoCA raw score for normal cognition (MoCA ≥26) and potential cognitive impairment (MoCA <26).

Multivariate linear regression was completed with SLIT2 plasma and vitreous protein level again utilized as a predictor of neurocognitive test score and adjusted for the single predictors of age, sex, race, diabetic status, diabetic retinopathy status, glaucoma, APOE genotype, and APOE risk category. APOE risk categories (Supplemental Table 1) were constructed based on population genotype prevalence patterns and associated likelihood of developing Alzheimer’s disease, as previously published.^43^ Multiple predictor analysis was also completed and adjusted for demographics, which included age, sex, and race, and for demographics with additional medical and ophthalmic history variables. A complete model, which included demographics, diabetic retinopathy status, glaucoma status, and APOE risk category, was also analyzed as applicable. Diabetes mellitus and diabetic retinopathy status could not be simultaneously included in the complete model due to variable collinearity. For all complete models, effect size was calculated to demonstrate the contribution of SLIT2 to the variation in neurocognitive test scores. Additionally, power, with an assumed effect size of 0.2 (α=0.05), was calculated to demonstrate the complete models’ ability to detect low-medium sized effects.

## RESULTS

### Demographics

Seventy-nine participants were enrolled in the study; 79 vitreous samples and 67 blood samples were collected. Missing samples were due to challenges with specimen collection, such as poor intravenous access, participant refusal of blood draws, hemolyzed blood, and inadequate sample volume. Adequate material for the SLIT2 assay was obtained in 69 vitreous and 60 plasma samples. MoCA testing was completed in 63 participants, and IRV testing was completed in 56. Figure 1 summarizes the biospecimen samples collected and the neurocognitive testing completed. Table 1 describes the study demographic characteristics.

**Figure 1.**
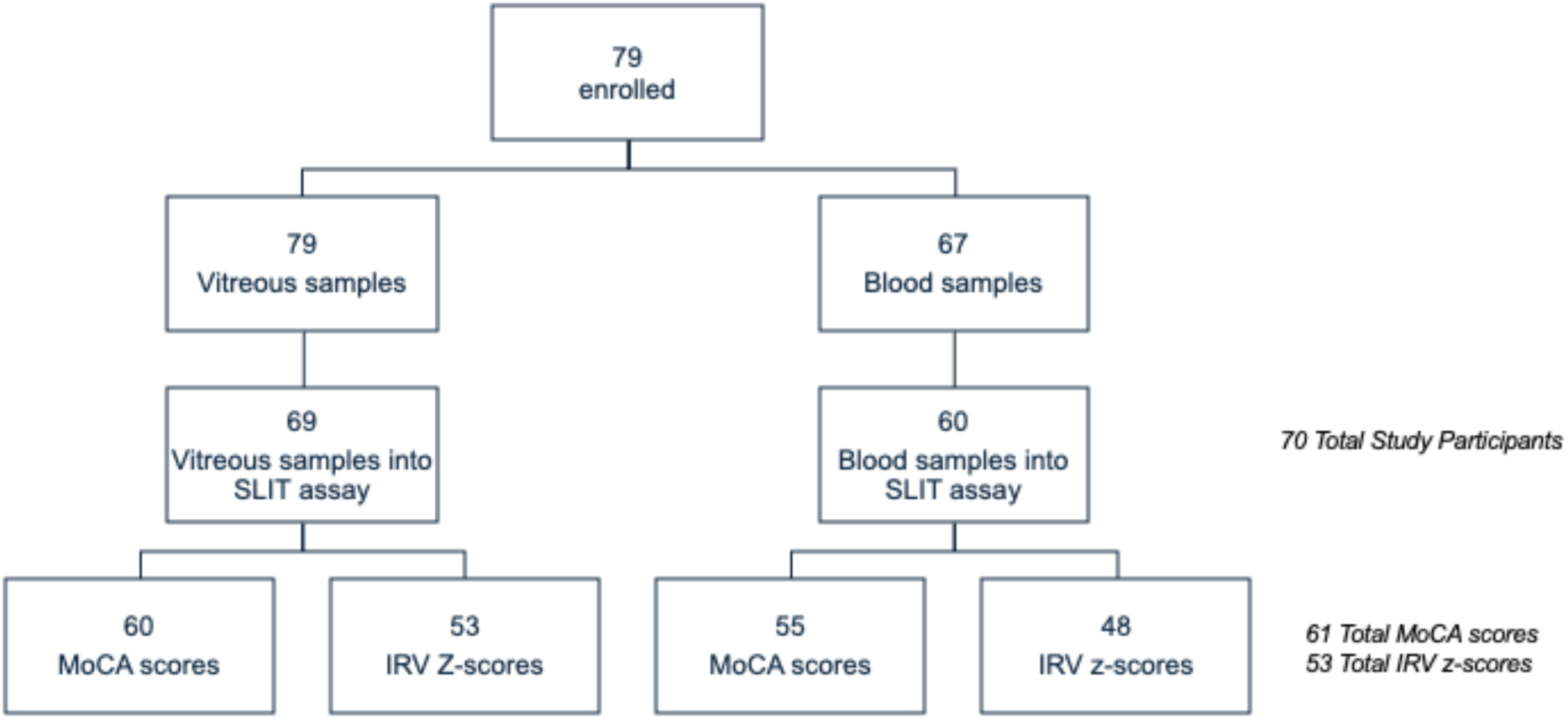
Enrolled Participant Flow Diagram. A total of 79 patients were enrolled in the study. Missing samples were due to challenges with specimen collection, such as poor intravenous access, participant refusal of blood draws, hemolyzed blood. Additionally, there was inadequate sample volume for SLIT assay analysis for some collected samples. Lastly, neurocognitive testing was not completed or unable to be obtained for all participants.

**Table 1:**
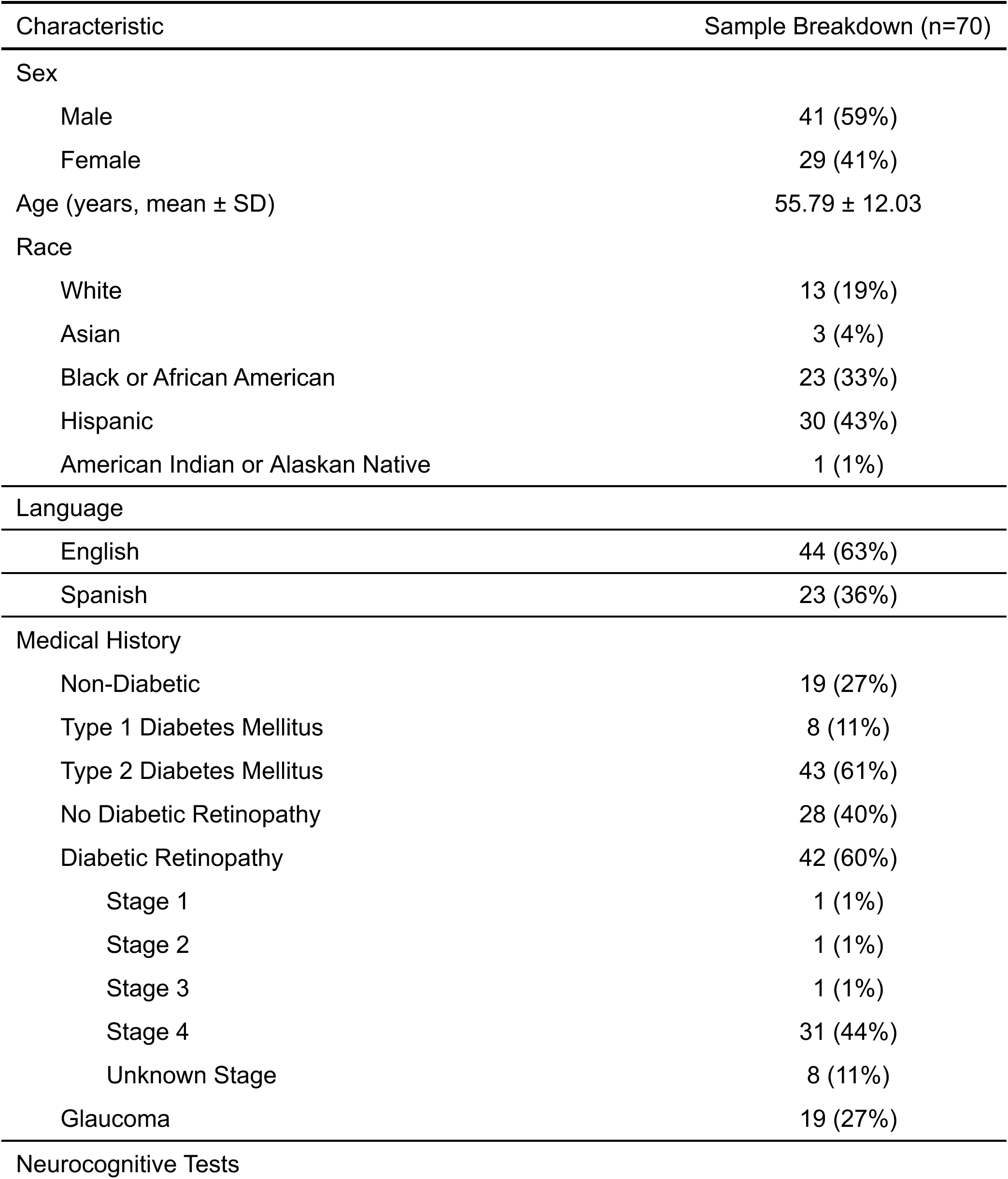
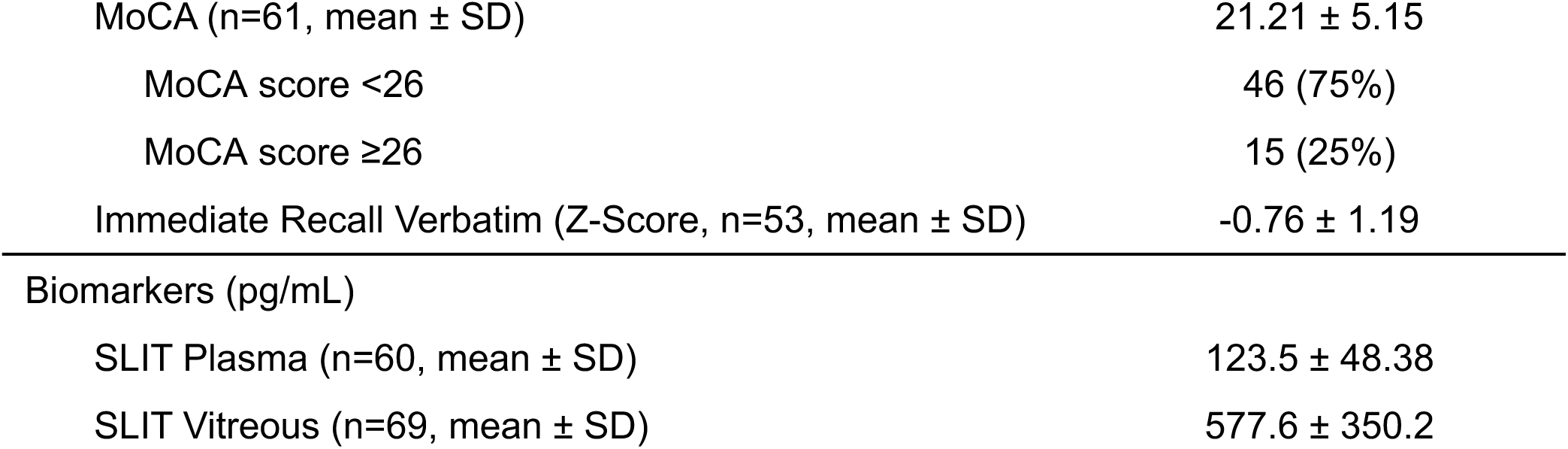
Participant Demographics and Medical History.

### SLIT2 Protein Detection

Our data showed that the newly developed highly sensitive SLIT2 MSD assay can detect SLIT2 protein expression levels as low as 5 pg/ml, which is 200-fold more sensitive than regular ELISA. In addition, we discovered a 7-fold higher SLIT2 level in the vitreous humor compared to blood plasma samples in humans. In a Spearman correlation, SLIT2 plasma and vitreous levels were not correlated with each other (*ρ*=-0.05424, p=0.6833).

### Univariate Analysis

In univariate linear regression, lower vitreous SLIT2 levels were associated with lower MoCA raw scores (m=0.003897, r^2^=0.07585, p=0.0332, Figure 2a) and lower IRV z-scores (m=0.001196, r^2^=0.09811, p=0.0224, Figure 2b). Alternatively, plasma levels showed the opposite relationship, with higher plasma SLIT2 associated with lower MoCA scores (m=-0.03559, r^2^=0.09699, p=0.0206, Figure 3a). Plasma SLIT2 levels were not associated with IRV z-scores (m=-0.003430, r^2^=0.01830, p=0.3593, Figure 3b).

**Figure 2.**
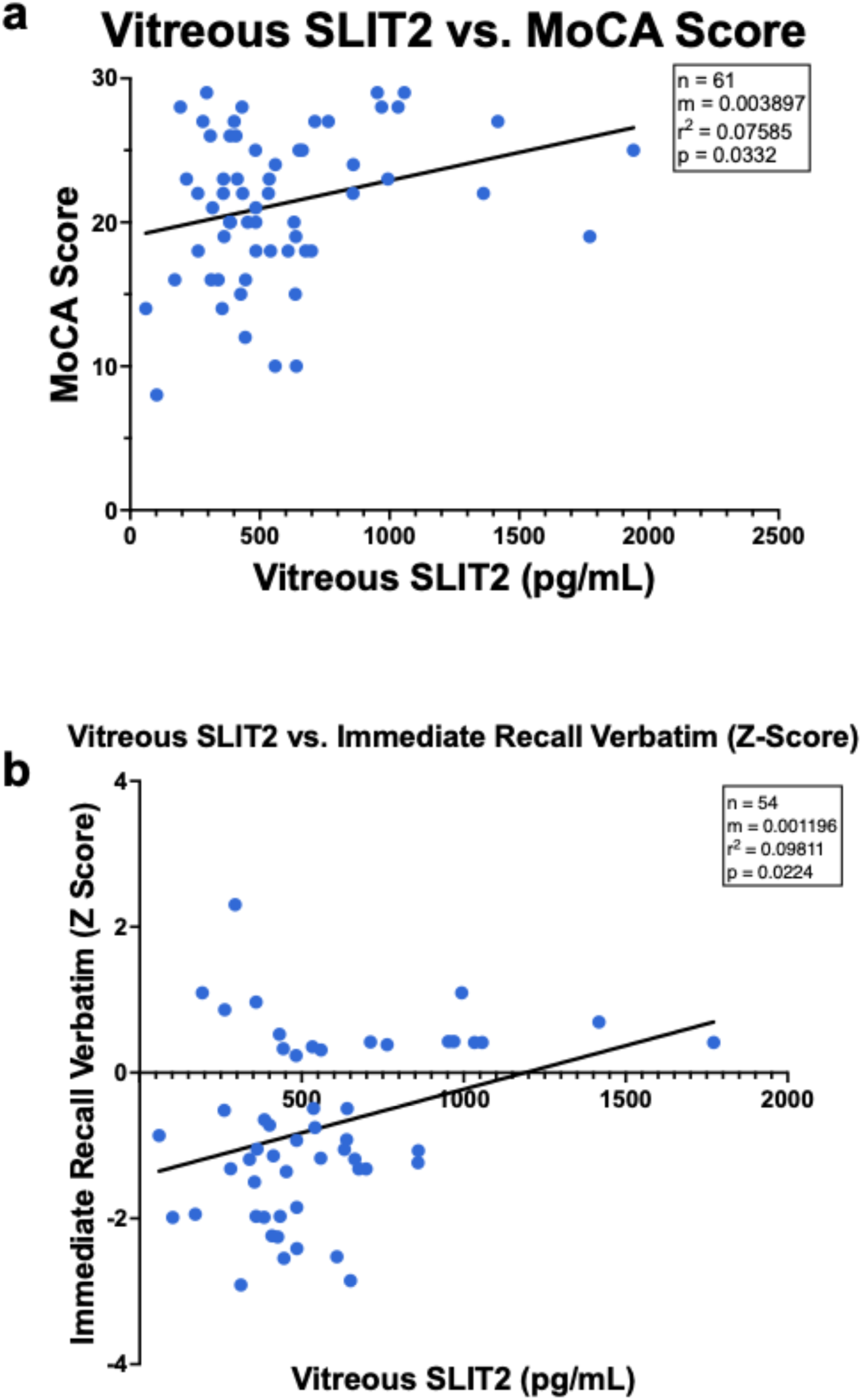
The relationship of SLIT2 in the vitreous with neurocognitive tests. Linear regression analysis demonstrated a significant relationship between SLIT2 vitreous protein level and MoCA score (a, r^2^=0.076, p=0.0332) and Immediate Recall Verbatim Z-Score (b, r^2^=0.098, p=0.0224).

**Figure 3.**
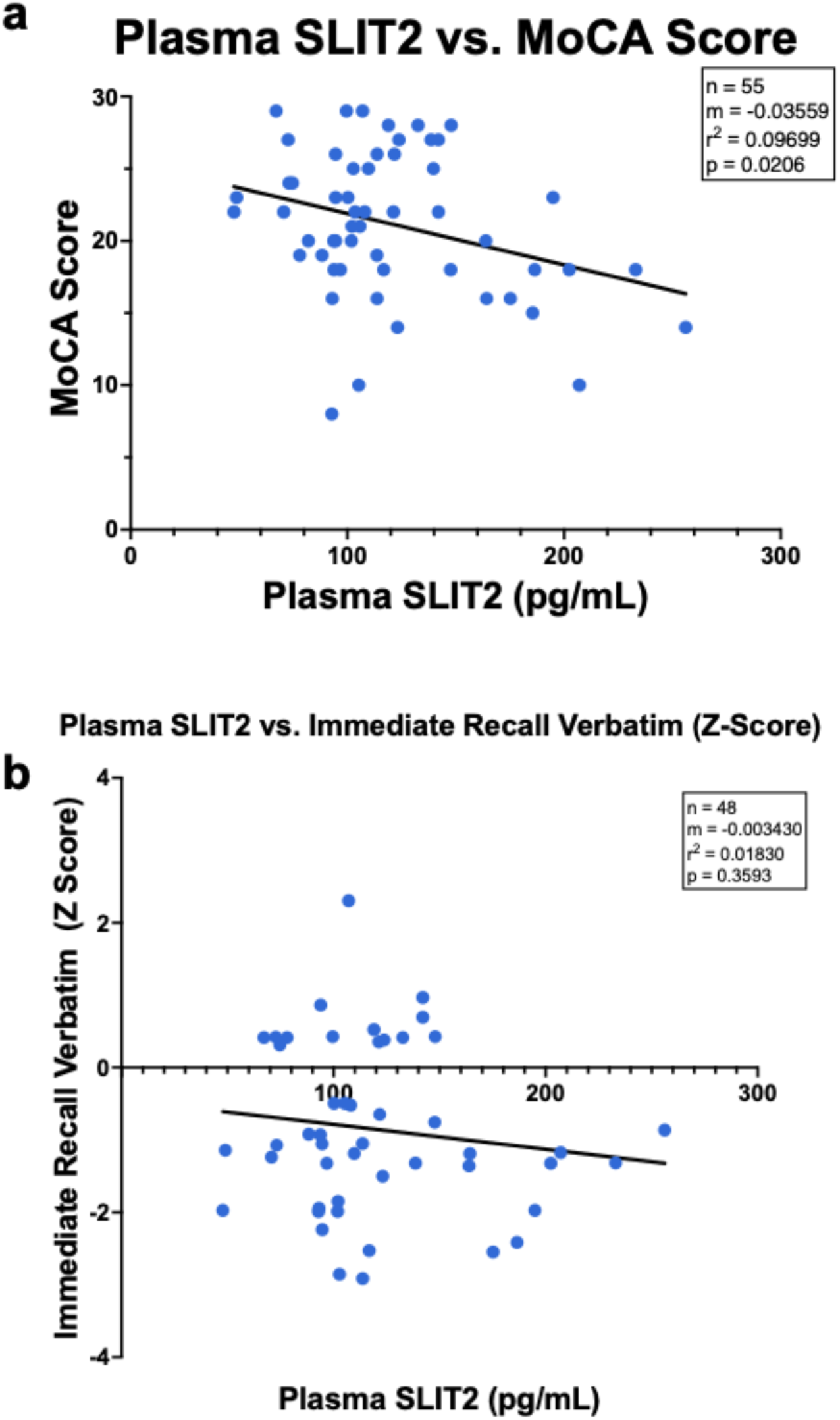
The relationship of SLIT2 plasma levels with neurocognitive tests. Linear regression analysis demonstrated a significant relationship between SLIT2 plasma protein levels and MoCA scores (a, r^2^=0.097, p=0.0206). There was no relationship between SLIT2 plasma protein levels and Immediate Recall Verbatim Z-scores (b, r^2^=0.01830, p=0.3593).

Further analysis among only individuals with a MoCA score indicating possible cognitive impairment (MoCA score below 26) was associated with lower vitreous SLIT2 (m=0.003660, r^2^=0.09902, p=0.0353, Figure 4a) and higher plasma SLIT2 (m=-0.03311, r^2^=0.0073, p=0.0073, Figure 4b). In individuals with a MoCA score in the normal cognition range (≥26), there was no significant association of MoCA scores with SLIT2 in plasma (m=-0.008733, r^2^=0.03803, p=0.5231, Supplemental Figure 2a) or SLIT2 in vitreous (m=0.003660, r^2^=0.09902, p=0.2789, Supplemental Figure 2b).

**Figure 4.**
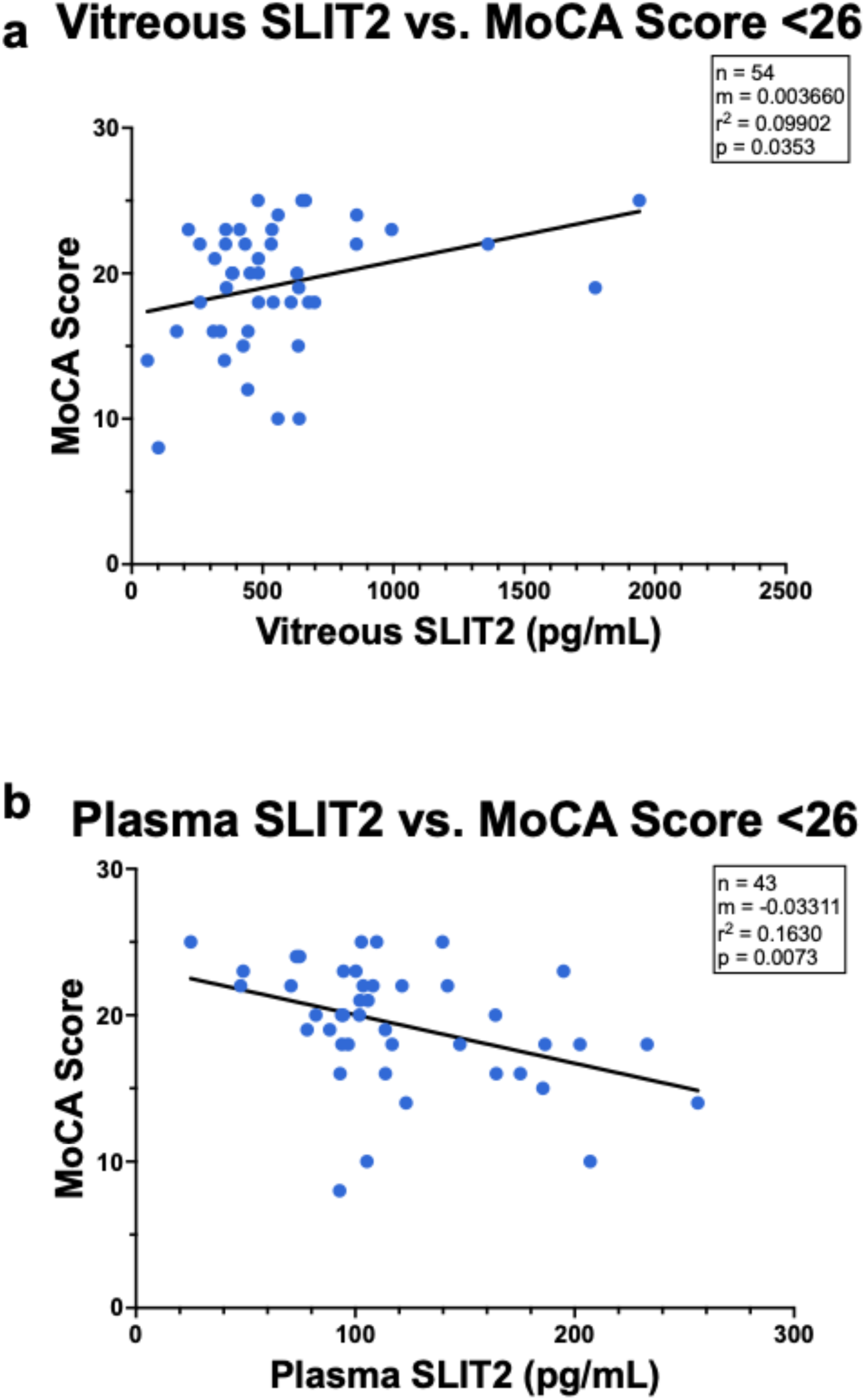
The relationship between SLIT2 and Montreal Cognitive Assessment (MoCA) Scores <26, the cut off value for mild cognitive impairment (MCI). Linear regression analysis demonstrated the relationship between SLIT2 vitreous protein level and MoCA scores <26 was significant (a, r^2^=0.099, p=0.0353). There was also a significant relationship between SLIT2 plasma protein level and MoCA scores <26 (b, r^2^=0.163, p=0.007).

### Multivariate Analysis

In single predictor adjustment models (Table 2), lower vitreous SLIT2 was associated with lower MoCA score after adjusting separately for age, sex, race, diabetic status, diabetic retinopathy status, and glaucoma. Multiple predictor models were significant when adjusted for demographics (age, sex, and race; p=0.0225), demographics with diabetic status (p=0.0162), demographics with diabetic retinopathy status (p=0.0269), and demographics with glaucoma status (p=0.0111). A complete model adjusting for demographics, diabetic retinopathy status, and glaucoma status was significant (p=0.0141) with a large effect size (f^2^=0.391) and sufficient power (92.5%, f^2^=0.2, α=0.05). APOE genotype and risk category were not included in the complete model for vitreous SLIT2 because the single predictor analysis for both was not significant.

**Table 2:**
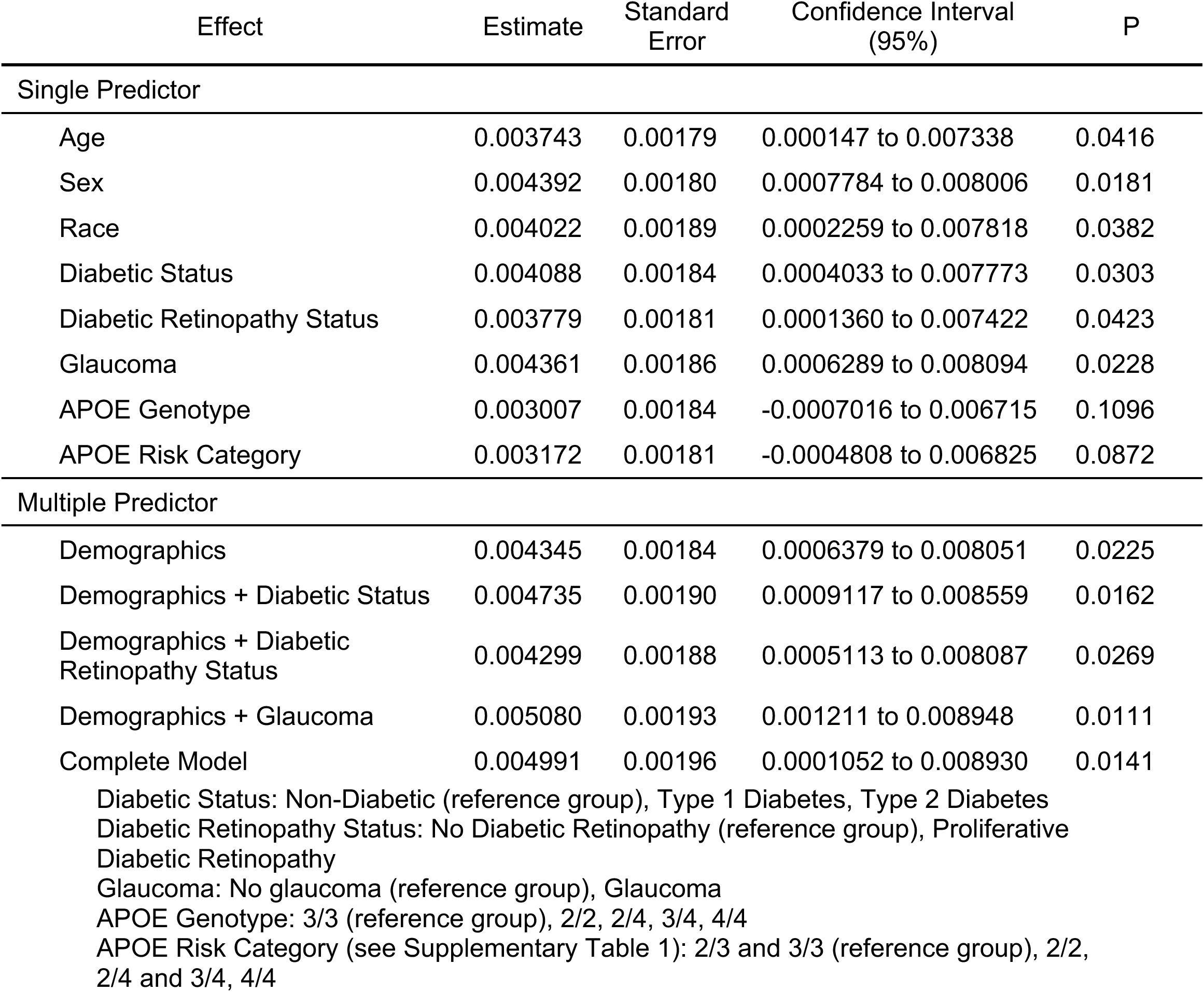
Predictor Effects on the Association of SLIT2 Vitreous and MoCA Score. The single predictor model applied each variable separately to determine its individual impact on the association of SLIT2 Vitreous and MoCA Score. In the multiple predictor model, all demographic variables (age, sex, and race) were additionally combined with medical variables. APOE genotype and risk category were not included in the multiple predictor or complete models because the single predictor model did not find a significant association. The complete model did meet statistical significance (p=0.0141) with a large effect size (f^2^=0.391) and sufficient power (92.5%, f^2^=0.2, α=0.05).

Higher plasma SLIT2 was associated with lower MoCA score in single predictor adjustment models adjusted separately for age, sex, race, diabetic status, diabetic retinopathy status, glaucoma, APOE genotype, and APOE risk category (Table 3). In multiple predictor analysis, similarly significant associations were found when adjusted for demographics (age, sex, and race; p=0.0498), demographics with diabetic retinopathy status (p=0.0198), demographics with APOE genotype (p=0.0418), and demographics with APOE risk category (p=0.0413). A complete model adjusting for demographics, diabetic retinopathy status, glaucoma status, and APOE risk category was not statistically significant (p=0.0522) but had a large effect size (f^2^=0.5026), signifying SLIT2 has a substantial contribution to the variation in MoCA score. In addition, we found sufficient statistical power (87.7%, f^2^=0.2, α=0.05) in the complete model.

**Table 3:** Predictor Effects on the Association of SLIT2 Plasma and MoCA Score. The single predictor model applied each variable separately to determine its individual impact on the association of SLIT2 plasma and MoCA Score. In the multiple predictor model, all demographic variables (age, sex, and race) were additionally combined with medical variables. For this analysis, APOE risk category was included in the complete model because the single predictor model was significantly associated with plasma SLIT2. The complete model did not meet statistical significance (p=0.0522) but had a large effect size (f^2^=0.5026) and sufficient power (87.7%, f^2^=0.2, α=0.05).

| Effect | Estimate | Standard Error | Confidence Interval (95%) | P |
| --- | --- | --- | --- | --- |
| Single Predictor |  |  |  |  |
| Age | -0.03469 | 0.01488 | -0.06454 to -0.004826 | 0.0237 |
| Sex | -0.03584 | 0.01504 | -0.06602 to -0.005657 | 0.0209 |
| Race | -0.03208 | 0.01562 | -0.06346 to -0.0007018 | 0.0453 |
| Diabetic Status | -0.03431 | 0.01551 | -0.06543 to -0.003179 | 0.0314 |
| Diabetic Retinopathy Status | -0.04446 | 0.01556 | -0.07569 to -0.01323 | 0.0061 |
| Glaucoma | -0.03749 | 0.01560 | -0.06882 to -0.006153 | 0.0200 |
| APOE Genotype | -0.03605 | 0.01559 | -0.06741 to -0.004692 | 0.0252 |
| APOE Risk Category | -0.03512 | 0.01555 | -0.06639 to -0.003847 | 0.0285 |
| Multiple Predictor |  |  |  |  |
| Demographics | -0.03096 | 0.01538 | -0.06190 to -0.0000256 | 0.0498 |
| Demographics + Diabetic Status | -0.03012 | 0.01612 | -0.06259 to 0.002351 | 0.0682 |
| Demographics + Diabetic Retinopathy Status | -0.03876 | 0.01605 | -0.97198 to -0.006437 | 0.0198 |
| Demographics + Glaucoma | -0.03266 | 0.01663 | -0.06622 to 0.0009003 | 0.0562 |
| Demographics + APOE Genotype | -0.03402 | 0.01619 | -0.06672 to -0.001326 | 0.0418 |
| Demographics + APOE Risk Category | -0.03375 | 0.01603 | -0.06610 to -0.001401 | 0.0413 |
| Complete Model | -0.03620 | 0.01803 | -0.07276 to 0.0003654 | 0.0522 |

Diabetic Status: Non-Diabetic (reference group), Type 1 Diabetes, Type 2 Diabetes Diabetic Retinopathy Status: No Diabetic Retinopathy (reference group), Proliferative Diabetic Retinopathy Glaucoma: No glaucoma (reference group), Glaucoma

Vitreous SLIT2 and IRV z-score were also significant in single predictor models adjusted separately for age, sex, race, diabetic status, diabetic retinopathy status, glaucoma status, and APOE risk category (Table 4), and in multiple predictor models adjusted for demographics (age, sex, and race; p=0.0142), demographics with diabetic status (p=0.0149), demographics with diabetic retinopathy status (p=0.0175), and demographics with glaucoma (p=0.0336). A complete model adjusting for demographics, diabetic retinopathy status, glaucoma status, and APOE risk category was not significant (p=0.1328) but had a large effect size (f^2^=0.568), signifying SLIT2 has a substantial contribution to variation in IRV z-score. This complete model was also found to be sufficiently powered (84.0%, f^2^=0.2, α=0.05).

**Table 4:** Predictor Effect on the Association of SLIT2 Vitreous and IRV z-score. The single predictor model applied each variable separately to determine its individual impact on the association of SLIT2 vitreous and IRV z-score. In the multiple predictor model, all demographic variables (age, sex, and race) were additionally combined with medical variables. The complete model did not meet statistical significance (p=1328) but had a large effect size (f^2^=0.568) and sufficient power (84.0%, f^2^=0.2, α=0.05).

| Effect | Estimate | Standard Error | Confidence Interval (95%) | P |
| --- | --- | --- | --- | --- |
| Single Predictor |  |  |  |  |
| Age | 0.00115 | 0.0004934 | 0.001588 to 0.002141 | 0.0239 |
| Sex | 0.001276 | 0.0005092 | 0.0002535 to 0.002299 | 0.0155 |
| Race | 0.001268 | 0.0005337 | 0.0001942 to 0.002342 | 0.0217 |
| Diabetic Status | 0.001249 | 0.0005158 | 0.002129 to 0.002286 | 0.0192 |
| Diabetic Retinopathy Status | 0.001155 | 0.0005102 | 0.0001306 to 0.002180 | 0.0279 |
| Glaucoma | 0.001107 | 0.0005086 | 0.00008515 to 0.002129 | 0.0343 |
| APOE Genotype | 0.001083 | 0.0005381 | -0.000004023 to 0.002169 | 0.0508 |
| APOE Risk Category | 0.001078 | 0.0005268 | 0.00001528 to 0.002141 | 0.0469 |
| Multiple Predictor |  |  |  |  |
| Demographics | 0.001332 | 0.0005220 | 0.0002807 to 0.002383 | 0.0142 |
| Demographics + Diabetic Status | 0.001345 | 0.0005353 | 0.0002755 to 0.002415 | 0.0149 |
| Demographics + Diabetic Retinopathy Status | 0.001300 | 0.0005266 | 0.0002387 to 0.002361 | 0.0175 |
| Demographics + Glaucoma | 0.001178 | 0.0005364 | 0.00009582 to 0.002259 | 0.0336 |
| Demographics + APOE Genotype | 0.001000 | 0.0006345 | -0.0002880 to 0.002288 | 0.1240 |
| Demographics + APOE Risk Category | 0.001053 | 0.0005828 | -0.0001287 to 0.002235 | 0.0791 |
| Complete Model | 0.0009285 | 0.0006025 | -0.0002972 to 0.002154 | 0.1328 |
| 639 Diabetic Status: Non-Diabetic (reference group), Type 1 Diabetes, Type 2 Diabetes<br>640 Diabetic Retinopathy Status: No Diabetic Retinopathy (reference group), Proliferative<br>641 Diabetic Retinopathy<br>642 Glaucoma: No glaucoma (reference group), Glaucoma<br>643 APOE Genotype: 3/3 (reference group), 2/2, 2/4, 3/4, 4/4<br>644 APOE Risk Category (see Supplementary Table 1): 2/3 and 3/3 (reference group), 2/2,<br>645 2/4 and 3/4, 4/4 |  |  |  |  |

## DISCUSSION

In this study, we found that vitreous and plasma SLIT2 protein were significantly associated with Montreal Cognitive Assessment (MoCA) and Immediate Recall Verbatim (IRV) scores. The relationship remained significant when adjusted for covariates of age, sex, race, diabetes, and diabetic retinopathy in single and multiple predictor models. We found up to a 7-fold increase in SLIT2 concentration in the vitreous humor compared to plasma. As far as we are aware, this is the first study to report relative concentrations of SLIT2 in vitreous and plasma and establish an association between SLIT2 levels from both sources with cognitive function obtained by MoCA and IRV z-score.

Our research partially supports previous studies that demonstrated associations between SLIT2 and cognitive function. But while a prior report found that higher plasma SLIT2 levels were associated with an increased risk of late-onset dementia,^36^ our study was conducted in a younger cohort of patients without known dementia diagnosis, indicating that our findings may be more relevant for MCI and early-onset dementia. There are no previous studies describing SLIT2 protein levels in early-onset dementia.

Additionally, our study further validates previous work done by our group that demonstrated an association between vitreous protein levels and neurocognitive test scores^19^ and the relatively higher neurodegenerative protein concentrations in the vitreous compared with plasma.^18^ We have also found in a prior post-mortem study that an elevated vitreous concentration may reflect an elevated cerebral concentration.^20^ In this study, our findings further demonstrate the potential for ocular fluids as a biospecimen sampling source for early detection and diagnosis of neurocognitive disorders, as well as the unexplored role of SLIT-ROBO signaling in the pathological development of neurodegenerative disease.

We found opposite directions of associations in the vitreous humor and plasma between SLIT2 levels and MoCA score, the reasons for which remain unclear. SLIT2 is a paracrine ligand associated with the extracellular membrane receptor ROBO, resulting in high concentrations in local tissues surrounding an area of increased expression. As SLIT2 is required for active axon pathfinding and new neuronal synapse formation in the central nervous system (CNS), one possibility is that SLIT2 is more highly expressed in the brain and ocular tissues compared to blood, and therefore lower SLIT2 in vitreous humor may suggest decreased axon pathfinding and fewer new neuronal synapse formations in the brain, observed as lower MoCA and IRV scores. Relatively higher plasma SLIT2 may indicate the result of a systemic body reaction attempting to increase SLIT2 levels and stimulate axon pathfinding and new neuronal synapse formations.

Similar inverse relationships have been noted in Aβ, with higher concentrations of amyloid plaque deposition in the brain being associated with lower Aβ concentrations in the CSF and plasma.^46–49^ As previously mentioned, our group has found significantly lower Aβ concentrations in the vitreous in individuals with lower cognitive scores (based on mini-mental status examination), consistent with what is typically noted in the cerebral spinal fluid (CSF) in those with MCI, early-onset dementia, and late-onset Alzheimer’s Disease.^11,19^ Additionally, some insight might be gathered from the inverted relationship expanding as neurocognitive function declines. It has been previously demonstrated that neurodegeneration is associated with impaired blood-brain barrier (BBB) and CNS vasculature functionality.^50^ Increased BBB permeability^51^ and decreased function in pathological inflammatory states may contribute to the increased concentration of SLIT2 in the plasma, as SLIT2 also plays a role in inflammation.^52,53^ The involvement of SLIT2 remains unclear, as SLIT2 overexpression in mouse models have been shown to have increased or decreased BBB permeability.^54,55^ Interestingly, overexpression of SLIT1, which is a homologue of SLIT2 and highly expressed in the brain, can facilitate functional neuronal regeneration in post-ischemic stroke mice by enhancing new neuron migration through the brain glial meshwork to stroke lesions.^56^ Further investigation into the mechanism by which SLIT2 exits the CNS and enters the systemic circulatory system is needed to better understand the relative concentration differences between vitreous and plasma samples.

SLIT2 was significantly associated with diabetes, which has been previously shown in rat models of type 1 diabetes^38^ and human epidemiological studies.^57–59^ Diabetes has been shown to be associated with higher rates of late-onset Alzheimer’s disease ^60^; elevated blood glucose, obesity, and metabolic disease have been shown to be potential risk factors for early-onset dementia.^12,61,62^ It is hypothesized that the pathophysiological mechanisms of hyperglycemia induce and increase the neuropathological changes.^60^

Limitations of this study include limited sample size and a lack of comprehensive neurological testing. The sample size may limit the precision of regression estimates, especially multivariate models with multiple predictors, increasing the potential for variability in sampling. However, we were able to achieve adequately powered complete models able to detect low-medium effects. While our sample was normally distributed with regard to MoCA score, IRV z-score, and SLIT2 protein level (Supplemental Figure 1), the limited numbers made it difficult to make more specific comparisons.

Additionally, further confirmation of MCI and dementia diagnosis, including potentially early-onset disease, in individuals with lower scores would require further testing and functional assessment of participants. The limited sample also prevented analysis adjusted for additional ocular co-morbidities, such as macular degeneration, which was not present in any of our participants, likely due to our younger and more racially and ethnically diverse study cohort. High prevalence of comorbidities known to be associated with dementia risk, including early-onset dementia risk^12,61,62^, such as diabetes, likely contributed to the low average neurocognitive score in our young study cohort. The large percentage of APOE E4 also biases our sample and reduces the generalizability of our findings. This may have contributed to the low average neurocognitive score, since it has been shown that diabetic patients are more genetically predisposed to late-onset Alzheimer’s Disease.^63^ Further prospective studies with larger recruitment goals may address these limitations. Additionally, participants for this study were patients from the eye clinic at Boston Medical Center and, therefore, were recruited based on ocular history, not neurological evaluation. While we adjusted for any significant covariates, such as diabetes mellitus, diabetic retinopathy, and glaucoma, the impact of eye and systemic diseases on SLIT2 levels remains unclear. Because of this study’s setting, only screening-level neuropsychological testing was performed. Further studies validating our findings on the basis of neurodegenerative diagnosis and staging could provide more specific information regarding the extent of neurodegeneration associated with SLIT2 protein levels.

In conclusion, we have demonstrated that SLIT2 protein in the vitreous and plasma is significantly associated with Montreal Cognitive Assessment score and Immediate Recall Verbatim z-score, and the relationship remained significant when adjusted for demographics, medical and ocular co-morbidities, and APOE genotype factors. The utilization of a custom-made highly sensitive MSD immunoassay allowed for quantification of SLIT2 at the pg/mL level. The inverse concentrations between vitreous and plasma samples highlight the need to further understand the mechanisms underlying neurodegeneration and its relationship with systemic circulation.

## DECLARATIONS

### Ethics Approval and Consent to Participate

BUMC/BMC institutional review board provided approval and oversight (study reference number H-37370, principal investigator MLS).

### Data Availability

The data supporting the findings of this study are available on request from the corresponding author.

## Author contributions

SIS (Conceptualization, Data Curation, Formal Analysis, Investigation, Methodology, Visualization, Writing – Original Draft); EL (Conceptualization, Investigation, Methodology, Writing – Original Draft); SA (Data Curation, Investigation, Project Administration, Writing – Review & Editing); ZS (Data Curation, Investigation, Project Administration, Writing – Review & Editing); SK (Data Curation, Investigation, Writing – Review & Editing); CH (Data Curation, Investigation, Writing – Review & Editing); IMS (Formal Analysis, Investigation, Methodology, Validation, Writing – Review & Editing); SN (Investigation, Writing – Review & Editing); XC (Investigation, Writing – Review & Editing); NHS (Investigation, Writing – Review & Editing); SSW (Conceptualization, Resources, Writing – Review & Editing); TS (Investigation, Methodology, Writing – Review & Editing); WL (Conceptualization, Data Curation, Formal Analysis, Funding Acquisition, Investigation, Methodology, Resources, Supervision, Writing – Original Draft, Writing – Review & Editing); MLS (Conceptualization, Data Curation, Formal Analysis, Funding Acquisition, Investigation, Methodology, Resources, Supervision, Writing – Original Draft, Writing – Review & Editing)

## Consent For Publication

Not applicable.

## Acknowledgments

We thank Dr. Weiming Xia and Dr. Michael Alosco of the Eye Biomarkers Study (EBS) and the BU Alzheimer’s disease center for their support. We thank Drs. Janice Weinberg and Jungwun Lee of the BU CTSI’s Biostatistics, Epidemiology & Research Design (BERD) program for their statistical consultation. We also thank Virginie Esain for providing technical support during the development of the SLIT2 MSD Assay.

## Supplemental Materials

**Supplemental Table 1:** APOE Allele Risk Categories.

| Risk Category | APOE Genotype | Plasma SLIT2 |  |  | Vitreous SLIT2 |  |  |
| --- | --- | --- | --- | --- | --- | --- | --- |
|  |  | n | Mean | SD | n | Mean | SD |
| 1 | 2/2 | 2 | 83.87 | 15.2 | 2 | 538.6 | 454.5 |
| 2 | 2/3<br>3/3 | 3 | 135.7 | 57.85 | 3 | 722.3 | 242.4 |
| 3 | 2/4<br>3/4 | 38 | 121.4 | 38.62 | 40 | 611.7 | 424.3 |
| 4 | 4/4 | 13 | 129.9 | 58.76 | 13 | 486.9 | 182.0 |

**Supplemental Figure 1.**
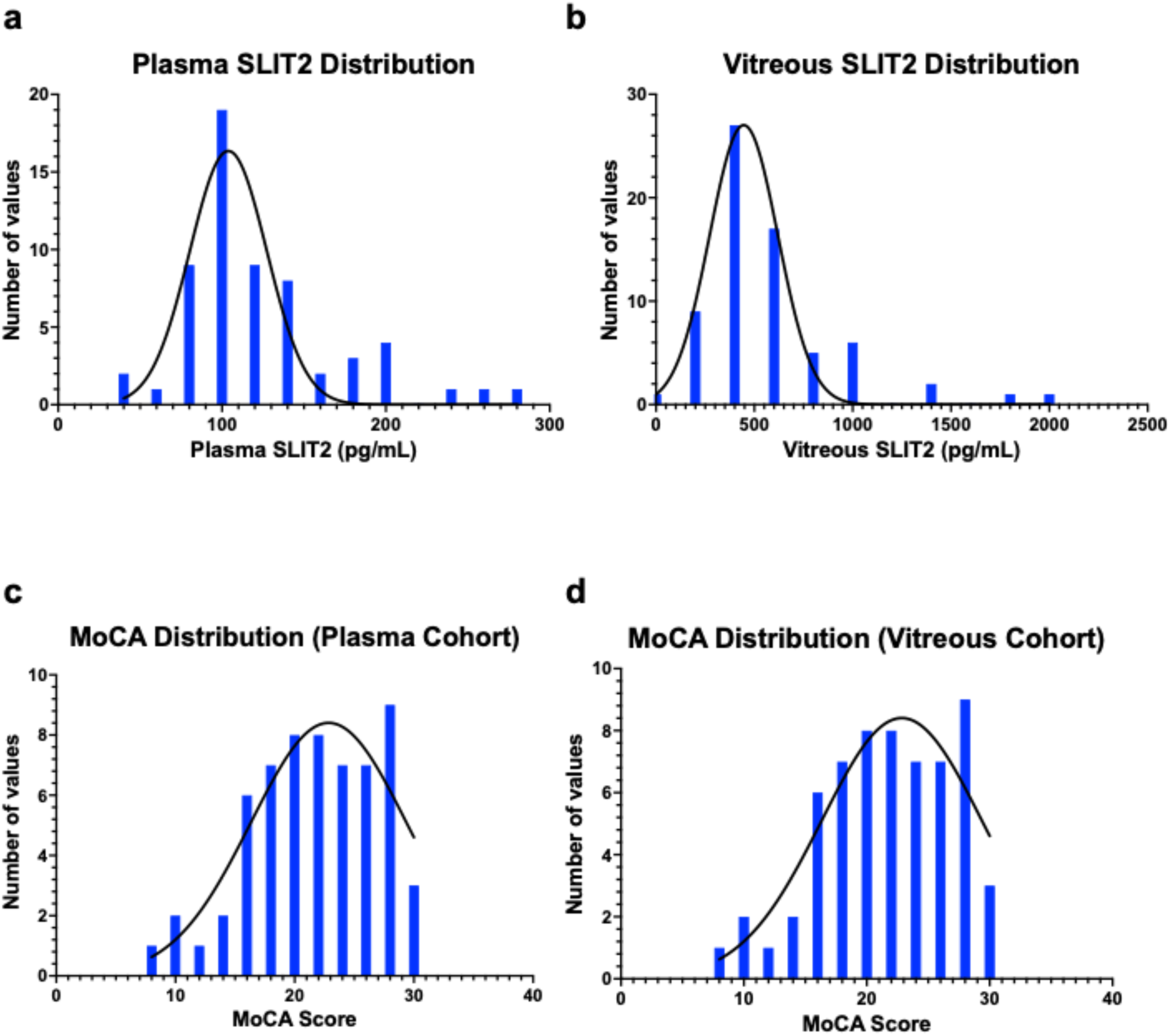
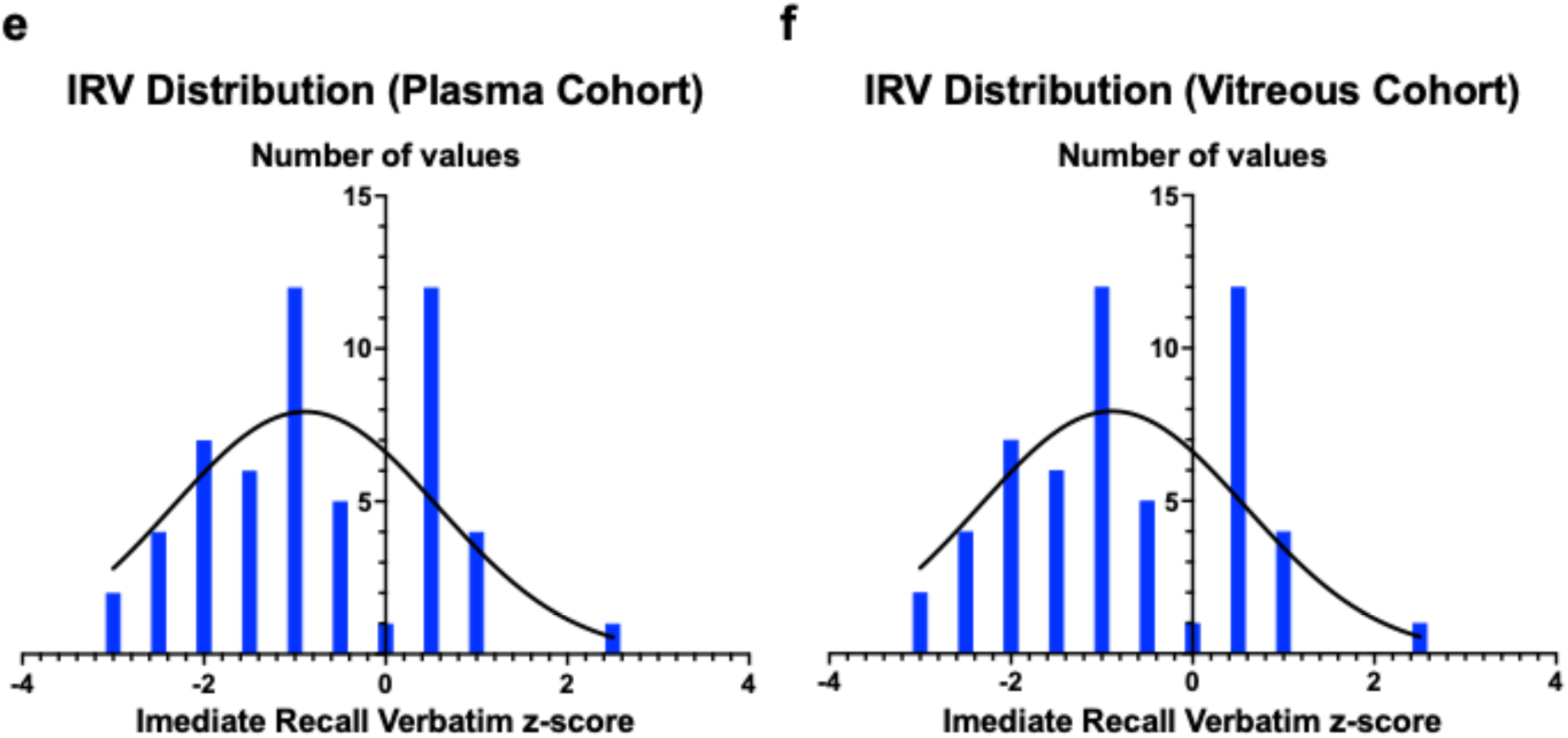
Distribution of SLIT2 protein levels and neurocognitive testing. Distribution of study predictor, SLIT2 plasma (a) and vitreous (b) level, and outcomes, Montreal Cognitive Assessment score (MoCA, c-d) and Immediate Recall Verbatim z-score (IRV, e-f) with overlying normalized curve.

**Supplemental Figure 2.**
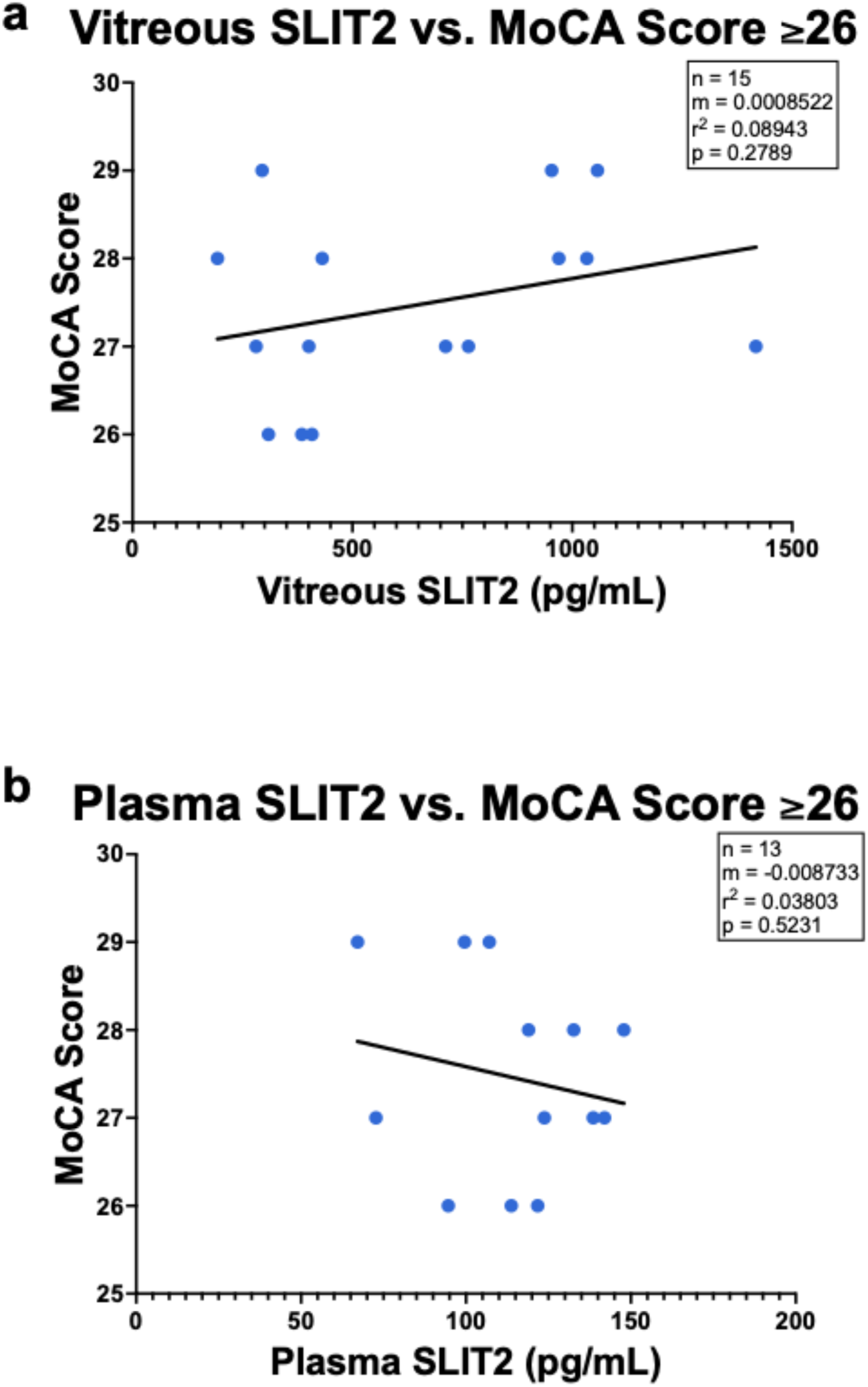
The relationship between SLIT2 and Montreal Cognitive Assessment (MoCA) Scores ≥26, the cut off value for mild cognitive impairment (MCI). Linear regression analysis demonstrated a moderately decreased relationship between SLIT2 protein in the plasma and MoCA scores >26 (a, r^2^=0.03803, p=0.5231) and a moderately increased relationship between SLIT2 protein in the vitreous and MoCA scores >26 (b, r^2^=0.08943, p=0.2789).

